# Enzyme stoichiometry indicates the variation of microbial nutrient requirements at different soil depths in subtropical forests

**DOI:** 10.1101/711069

**Authors:** Jiebao Liu, Ji Chen, Guangshui Chen, Jianfen Guo, Yiqing Li

## Abstract

Soil extracellular enzyme activities and associated enzymatic stoichiometry are considered sensitive indicators of nutrient availability and microbial substrate limitation. However, many of previous studies have been focusing on uppermost soil layer with a single enzyme as representative of the whole nutrient acquisition, leading to critical uncertainties in understanding soil nutrient availability and its relationship with microbial activities in deeper soils. In the current study, we investigated C-, N- and P-acquiring enzyme activities across a range of soil layers (0 - 10, 10 - 20, 20 - 40 and 40 - 60 cm), and examined the microbial C, N and P limitation in natural secondary forests (NSF) and Chinese fir (*Cunninghamia lanceolata*) plantation forests (CPF) in subtropical China. The results showed that microbial C and P co-limitation was detected in the two typical subtropical forests at all soil depths, rather than microbial N limitation. Microbial C and P limitation fluctuated along soil depth, but higher N was demanded by microbes in soil under 20 cm in both forests. The present results highlight the asymmetrical patterns of microbial nutrient limitation along the whole soil profile, and provide essential information in understanding nutrient limitations in deeper soils. These vertical and asymmetrical nutrient limitation patterns should be incorporated into future research studies priority.

## 1 Introduction

Tropical and subtropical forests possess rapid growth rates compared with their temperate counterparts and on a global scale, are more productive and sequester more atmospheric CO_2_ [1–3]. Tropical forest ecosystems are generally phosphorus (P) limited and at high altitudes are nitrogen (N) limited as well [4, 5]. The storage and subsequent release of P and N from soil for plant use is primarily governed by the action of soil micro-organisms. However, microbial processes are also limited or co-limited by carbon (C) (carbon can also be considered as energy source) or key nutrients (usually N or P) [6]. Therefore, determining soil microbial nutrient limitation is critical to understanding the nutrient availabilities and limitations of the tropical and subtropical forest ecosystems.

Soil enzymes are the key drivers of nutrient cycling in soils and are produced by both microbes and plants as root exudates. These enzymes are responsible for decomposition, turnover and mineralization of soil organic matter. More importantly, enzyme relative abundance or their stoichiometry reflects nutrient stoichiometric demands of microorganisms and environmental nutrient availability [7–11]. These enzymes include those that involved in the decomposition of cellulose (β-glucosidase (BG), cellobiosidase (CBH)), xylane (β-xylosidase, BX), chitin (β - 1,4-N-acetylglucosaminidase, NAG), polypeptides (leucine aminopeptidase, LAP) and phosphate (acid or alkaline phosphatase, AP) [12–15]. In particular, soil enzymes can be associated with mineral and organic matter as well as mineral-organic aggregates after release, which will alter enzymatic efficiencies due to the accessibility limitation of substrates. Therefore, recalcitrant substrates will induce microbes to secrete specific enzyme types and this can be additionally altered by product-substrate feedback [16]. However, enzymes involved in C and N mobilization in forest soils are usually represented by a single hydrolytic enzyme such as β-1,4-glucosidase (BG) and β-1,4-N-acetylglucosaminidase (NAG), respectively [17]. A previous meta-analyses have concluded that microbial C:N:P acquisition ratios converged on a 1:1:1 scale and this has hinged on the activities of ln(BG): ln(LAP + NAG): ln(AP) [17]. In general, degradation of organic matter requires the interaction of numerous enzyme types and using a single enzyme activity as an indicator of nutrient dynamics in soil is not an accurate reflection of the complexity of the forest ecosystem and most likely misrepresents metabolic activities [18–20].

Recently, the use of enzyme stoichiometry for assessing nutrient cycling has been suggested as an efficient method to indicate the relative resource limitation of soil micro-organisms. The theory of enzymatic stoichiometry has been widely used in different ecosystems [4, 17, 21–24], and a new vector analysis of eco-enzyme activities based on Sinsabaugh et al. (2008) [17] has been proposed by Moorhead et al. (2013, 2016) [6, 25]. The vector analysis plotted the proportion of C/N versus C/P acquiring enzyme activities, and then calculated an enzyme activity vector as the distance (length) and angle from the origin, finally quantify relative C vs. nutrient acquisition and the relative P vs. N acquisition. Based on stoichiometric and metabolic theories of ecosystems, increasing vector length was interpreted as a relative increase in C limitation and increasing vector angle as a relative increase in P vs. N limitation [25].

Vector analysis had been used to assist in understanding which resources limit microbial growth and activity [23, 26–28]. However, studies on enzyme activities together with enzymatic stoichiometry utilizing whole C-acquiring enzymes as a group are rare in forest ecosystems. Besides, the relative shortage of microbial nutrient limitation studies in subtropical forest ecosystem constrains our ability to accurately predict how soil nutrient cycle in forests will respond to climate change.

A large-scale study in north-south transect of Eastern China (NSTEC) with a unique vegetation belt ranging from boreal forest to tropical rain forest reported that microbial nutrient acquisition was P-limited in tropical forests [29]. However, other studies have indicated that karst and non-karst forests in southwest China were P-limited and microbial nutrient acquisition in karst ecosystems was more C- and P-limited rather than N-limited [23, 30]. Additionally, atmospheric N deposition will aggravate microbial C-limitation [31]. There have been few studies that examined enzymatic stoichiometry in fast-growing forest ecosystems of coniferous Chinese fir plantations (*Cunninghamia lanceolata*) (CPF) although these play key roles in carbon sequestration.

The most current enzymatic stoichiometry studies focused on topsoil (0-20 cm) [17, 23, 29, 30]. However, soils below 20 cm contribute over 50% of the global soil carbon stocks despite their low carbon concentration and their stability is questionable under different land use change and future global warming [32, 33]. The limited studies to date have found no consistent behaviors of soil enzymes and enzymatic stoichiometry in subsoil [14, 21, 34–36]. In addition, substrate limitation in subsoil is a major factor limiting microbial activity and controlling C turnover [37, 38]. Hence, exploring enzymatic stoichiometry patterns together with microbial nutrient limitation in subsoil can provide insights of biological mechanisms on nutrition cycling.

We hypothesized that microbial C, N and P acquisition would be higher in the fast-growing CPF than in the other forest due to its more rapid growth. Evergreen broadleaf forests and coniferous plantation forests are the main forest types in subtropical China. Therefore, paired natural secondary forests (NSF) and fast-growing coniferous Chinese fir plantation forests (CPF) were selected in the study to examine soil enzyme activities, enzymatic stoichiometry and microbial C, N and P limitation along the whole soil profile. Specifically, our objectives here were to (1) measure microbial C, N and P limitation across soil profile in both subtropical forests and (2) compare the effects of different calculation of C-acquiring enzymes on results of microbial nutrient limitation.

## 2 Materials and methods

The field studies did not involve endangered or protected species, and no animals were killed specifically for this study. All of our field studies and sampling procedures were with permission and the help of the Forest Bureau of Shunchang County.

The study was conducted in Shunchang County (117° 29’ ~ 118° 14’ E, 26° 38’ ~ 27° 12’ N) located in northwest Fujian province, China. The climate in the study area is subtropical marine monsoon with the annual mean temperature of 19.0 °C and the mean annual precipitation of 1629 mm. The soil at the sites is classified as red soil based on Chinese Soil Classification System [39], equivalent to clay loam Ferric Acrisol [40]. Both natural secondary forests (NSF) and coniferous Chinese fir plantation forests (CPF) used in the study that originated from the same natural evergreen forests at a 250 meter elevation. Three blocks were selected with a distance of approximately 3 km from each other (Fig. 1). In each block two plots were established, one in plantation and one in secondary forest, with a plot size of 20 × 20 m. The distance between the paired plots was about 50 m. Under the warm and moist climate, soils generally had no organic layer or the thickness of organic layer was no more than 1 cm. In this study we focused on mineral soils. The soil profile was divided into four layers: 0-10 cm, 10-20 cm, 20-40 cm, and 40-60 cm.

**Fig. 1.**
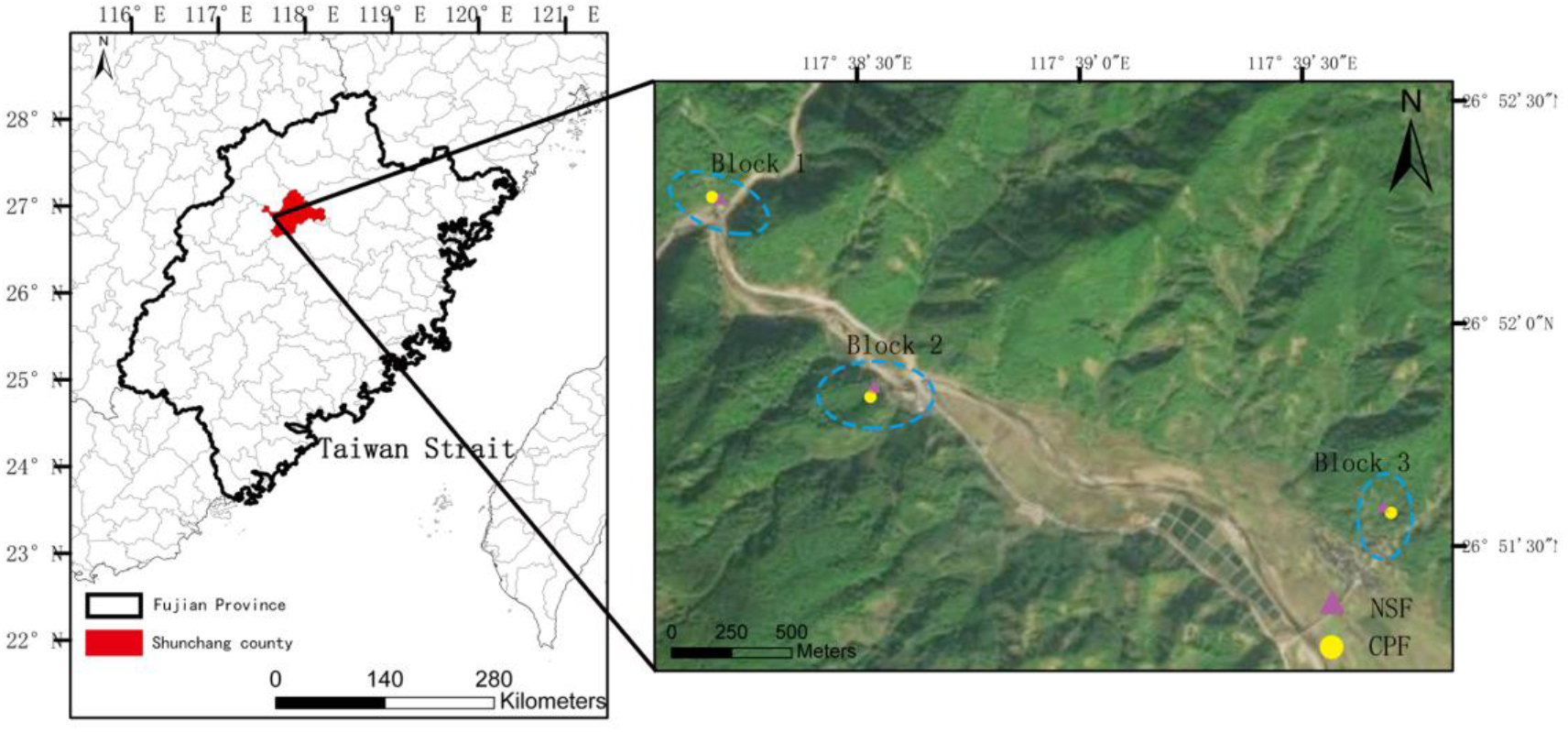
Study site locations of the natural secondary forest (NSF) and the coniferous fir plantation forest (CPF)

The dominant tree species in NSF are *Castanopsis fargesii* Franch, *Castanopsis lamontii*, *Castanopsis fargesii*, *Altingia gracilipes*. The main understory plant species in NSF included *Machilus chrysotricha*, *Woodwardia japonica*, *Adinandra millettii*, *Ardisia crispa* (Thunb.), *Sarcandra glabra* (Thunb.) and *Embelia rudis*. Canopy coverage is approximately 75%, the mean tree height and stand density were 14.7 m and 1267 stem ha^−1^, respectively (S1A Fig).

The CPF is dominated by Chinese fir (*C. lanceolata*) and dominant understory species are *Machilus grijsii* Hance, *Vaccinium bracteatum* Thunb, *Cleredendrum cwtophyllum* Turcz, *Dicranopteris dichotoma* Thunb. and *Rhizoma Cibotii*. Canopy coverage, mean tree height and stand density were 90%, 18.6 m and 1725 stem ha^−1^, respectively (S1B Fig).

### 2.1 Soil sampling

Soil samples were taken in August 2017 and after removal of surface debris, soil samples were collected at four soil layers (0 - 10, 10 - 20, 20- 40 and 40 - 60 cm) from five random locations in each plot. In each sampling plot, five soil cores at each layer were taken and mixed to represent a composite sample. Samples were immediately placed in sealed cooler containers and transported to the laboratory for chemical analysis. Stones and root fragments were removed and soils were sieved with a mesh size of 2 mm before chemical analyses. Soil physiochemical properties and nutrient stoichiometry are shown in Table 1.

**Table 1.**
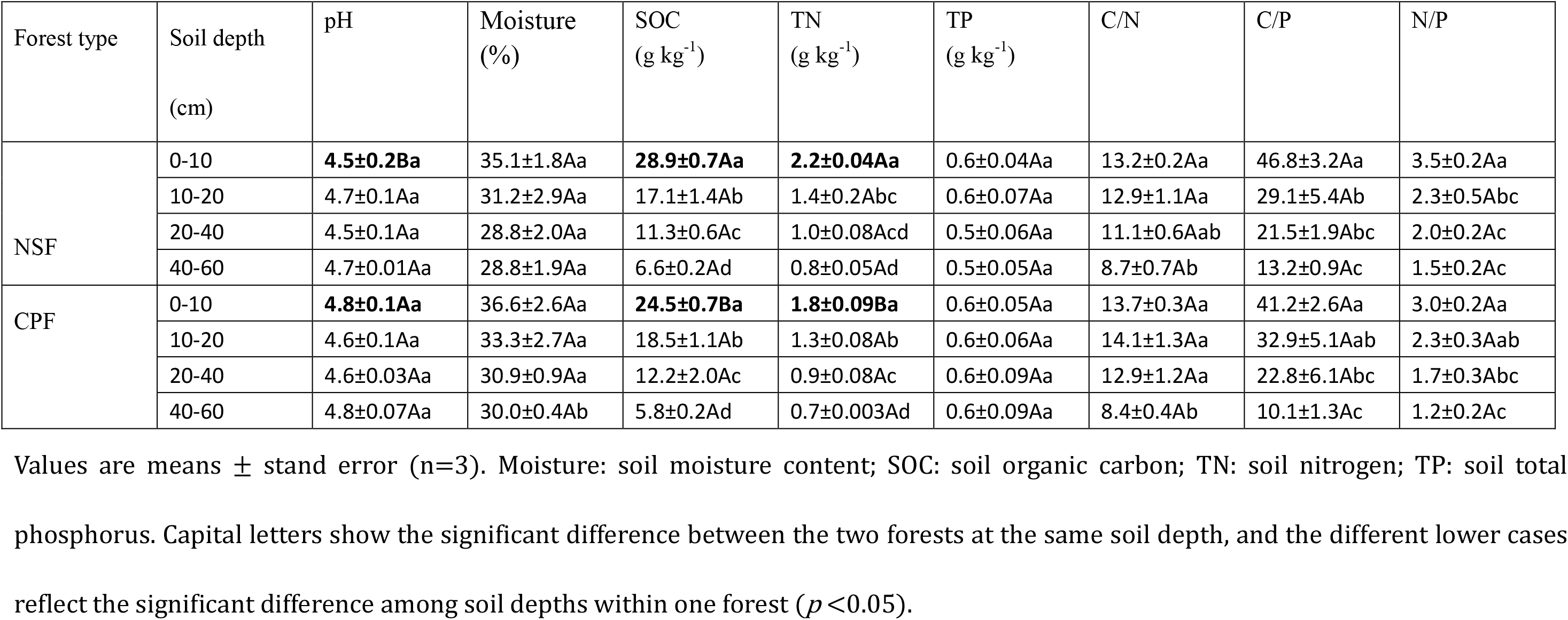
Soil physiochemical properties from the natural secondary forest (NSF) and Chinese fir plantation forest (CPF) at different soil depths.

### 2.2 Soil analysis

Soil moisture content was measured by drying 5 g soil at 105 °C for 24 h to a constant weight. Soil pH was determined using a pH meter with a ratio of water-to-soil at 2.5:1 (v/w). Soil organic carbon (SOC) and total nitrogen (TN) were determined by dry combustion in a Vario MAX CN elemental analyzer (Elementar Vario EL III, Germany). Soil total phosphorus (TP) was assayed using a Continuous Flow Analytic System (Skalar, The Netherlands).

Extracellular enzyme activities were determined following previously published methods [41, 42]. Fluorogenic methylumbelliferone-based (MUB) or methylcoumarin-based (MCA) artificial substrates were used to estimate the activities of C-cycling xylanases (BX), cellobiohydrolase (CBH), β-glucosidase (BG), N-cycling β −1,4-N-acetylglucosaminidase (NAG), peptide degrading leucine aminopeptidase (LAP) and P-cycling acid phosphatase (AP). High substrate concentration of each enzyme was adopted to ensure each enzyme to be assayed under saturation conditions. Briefly, suspensions of 1 g soil (dry weight equivalent) with 125 mL of 50 mM acetate buffer (pH 5.0) were dispersed using low-energy sonication (50 J s^−1^ output energy) [43]. The soil slurries and substrate solutions were incubated for 180 min at 20 ± 0.5 °C. Fluorescence was measured in microplates at an excitation wavelength of 365 nm and an emission wavelength of 450 nm with a Synergy H4 Hybrid Microplate Reader (Biotek, Winooski, VT, USA). To ensure that all enzyme activities were comparable, consistent parameters of incubation method and quenching were performed [8]. The incubation plates were shaken at an interval of once per hour to ensure reaction completion. The maximum duration of keeping soil slurries in buffer was less than 30 min before adding MUB (MCA)-linked substrate [44]. Fluorescence was conducted 1 min following the addition of 10 μl 1.0 M NaOH solution [42]. The time between the addition of substrate and the fluorescence reading was standardized to 180 min.

Units for enzyme activity were expressed as nmol activity g^−1^ dry soil h^−1^. C acquisition was expressed by the sum of BX, CBH and BG activities. N acquisition was represented by the sum of NAG and LAP while P acquisition was indicated as AP activity. The microbial C/N acquisition ratio was calculated as sums of hydrolytic enzymes using (BG + BX + CBH)/ (NAG + LAP), while (BG + BX + CBH)/ (AP) and (NAG + LAP)/ (AP) were used for the microbial C/P and N/P acquisition ratios, respectively.

### 2.3 Analysis of enzymatic stoichiometry

The following three approaches were used to examine microbial resource acquisition. The first was based on the method of Hill [22] to draw scatter plot of enzymatic stoichiometry, with Enzyme N/P as x axis and Enzyme C/ N as y axis. Due to the deviation from expected enzyme N/P (1:1) or enzyme C/N (1:1) ratio [17], different resource constraints (N limitation, P limitation, C&P limitation and N&P limitation) were shown in four district respectively. The second approach was based on enzyme ratios of C/P and N/P. Lower enzyme C/P and N/P ratios are suggestive of a degree of P limitation [4]. The third approach was vector analysis as previously reported [6].

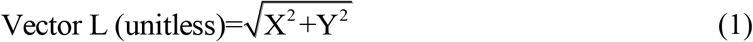

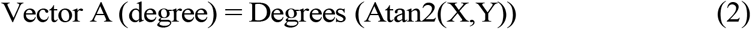

Where

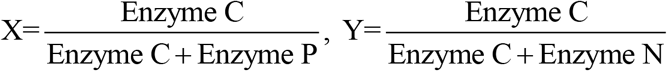

A relatively longer vector L (length) indicates greater C- acquisition, and the vector A (Angle) <45° and >45° indicate relative degrees of N- and P-limitation, respectively [26, 30].

### 2.4 Statistical analysis

One-way ANOVA followed by LSD test was used to determine the significance of enzyme activities and soil enzyme stoichiometry among the soil depths. Natural-log data transformations were used to meet the assumption of normality and homoscedasticity. All statistical analyses were done by SPSS v 23.0 (IBM SPSS Statistics for Windows, ver. 23.0; IBM, Armonk, NY, USA). Mixed linear model analyses in SPSS were selected to determine the effects of forest type, soil depth and their interactions on soil enzyme activities and enzymatic stoichiometry. Canonical correspondence analysis (CCA) was performed by using Canoco 5.0 software to test what environmental factors drive the enzymatic activities at whole soil depth.

## 3 Results

### 3.1 Soil enzyme activity and enzymatic stoichiometry

Results from mixed linear model showed that soil depth significantly influenced Enzyme C, N and P acquisitions, while the interactions of forest type × soil depth significantly affected enzyme C (Table 2; *p*<0.05). From the soil depths of 0-10 cm to 40-60 cm, the mean activities of C, N, and P acquisition enzymes in NSF and CPF decreased by 64%, 48%, and 76%, respectively (Fig. 2). We found no significant differences for C- acquiring (BX+ CBH+ BG) and P-acquiring (AP) enzyme activities between NSF and CPF at any soil layer. The only significant difference we found between the 2 forests was for N-acquiring (NAG + LAP) enzyme activities at surface soil (0-10 cm) and NSF activities were significantly greater than CPF (Fig. 2).

**Table 2.** Estimated fixed effects of soil depth, forest type and their interactions on soil enzyme, enzyme stoichiometry.

| Parameters | Intercept |  |  | Forest type |  |  | Soil depth |  |  | Forest type × Soil depth |  |  |
| --- | --- | --- | --- | --- | --- | --- | --- | --- | --- | --- | --- | --- |
|  | DF | F | *P* | DF | F | *P* | DF | F | *P* | DF | F | *P* |
| Enzyme C | 1 | 167.91 | < **0.001** | 1 | 0.02 | 0.90 | 3 | 26.40 | < **0.001** | 3 | 4.45 | < **0.05** |
| Enzyme N | 1 | 989.04 | < **0.001** | 1 | 6.63 | 0.06 | 3 | 34.74 | < **0.001** | 3 | 2.34 | 0.13 |
| Enzyme P | 1 | 179.17 | < **0.001** | 1 | 1.58 | 0.28 | 3 | 66.23 | < **0.001** | 3 | 2.30 | 0.15 |
| Enzyme C/N | 1 | 164.98 | < **0.001** | 1 | 1.75 | 0.27 | 3 | 3.69 | 0.07 | 3 | 2.17 | 0.18 |
| Enzyme C/P | 1 | 810.04 | < **0.001** | 1 | 10.04 | < **0.05** | 3 | 15.76 | < **0.001** | 3 | 5.91 | < **0.05** |
| Enzyme N/P | 1 | 197.25 | < **0.001** | 1 | 0.75 | 0.43 | 3 | 40.82 | < **0.001** | 3 | 3.13 | 0.08 |

C acquisition enzyme was represented by the sum of β-1,4-glucosidase (BG), cellobiohydrolase (CBH), β-xylosidase (BX). The sum of β-1,4-N-acetylglucosaminidase (NAG) and leucine aminopeptidase (LAP) were used for N acquisition. Acid phosphatase (AP) represented P acquisition enzyme. Enzymatic stoichiometry of C/N, C/P and N/P was calculated by the C, N and P acquisition enzyme respectively. Significant results ( $p < 0.05$ ) are indicated in bold.

**Fig. 2.**
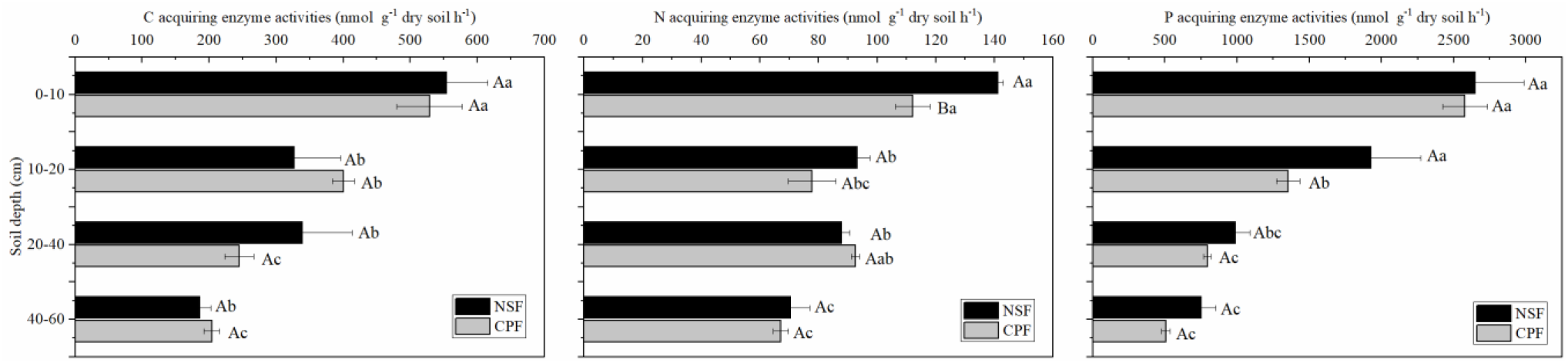
Variations of C, N and P acquiring enzyme activities at different soil depths of the natural secondary forest (NSF) and Chinese fir plantation forest (CPF). Statistically significant differences were assumed at *p* <0.05. All values are presented as mean ± standard error (n=3). Capital letters show the significant difference between the two forests at the same soil depth, and the different lower cases reflect the significant difference between soil depths within one forest.

In the NSF, the C- and N-acquiring enzyme activities displayed similar trends in their distributions. The surface soil layer had significantly greater enzyme activity than at the other 3 sampling layers (Fig. 2). In addition, there were no significant differences in P-acquiring enzyme activity at the 0-10 and 10-20 cm soil layers but both of these levels were significantly higher than at 20-40 and 40-60 cm (Fig. 2).

In the CPF, both C- and N-acquiring enzyme activities in the surface soil layer were significantly higher than at the lower levels (Fig. 2). Enzymatic P activity at 0-10 cm was almost 2-fold greater than at 10-20 cm and significantly higher than at 20-40 and 40-60 cm (Fig. 1).

The activity ratio of C/P was significantly affected by soil depth, forest type and their interactions, while the N/P ratio was only affected significantly by soil depth (Table 2, *p*<0.05). From 0–10 cm to 40-60 cm soil depth, mean soil enzymatic ratios of C/P and N/P increased by 59% and 119%, respectively (Fig. 3).Throughout the study area, the mean soil ratios of C/N, C/P and N/P were 3.88, 0.27 and 0.08, respectively (Fig. 3).

**Fig. 3.**
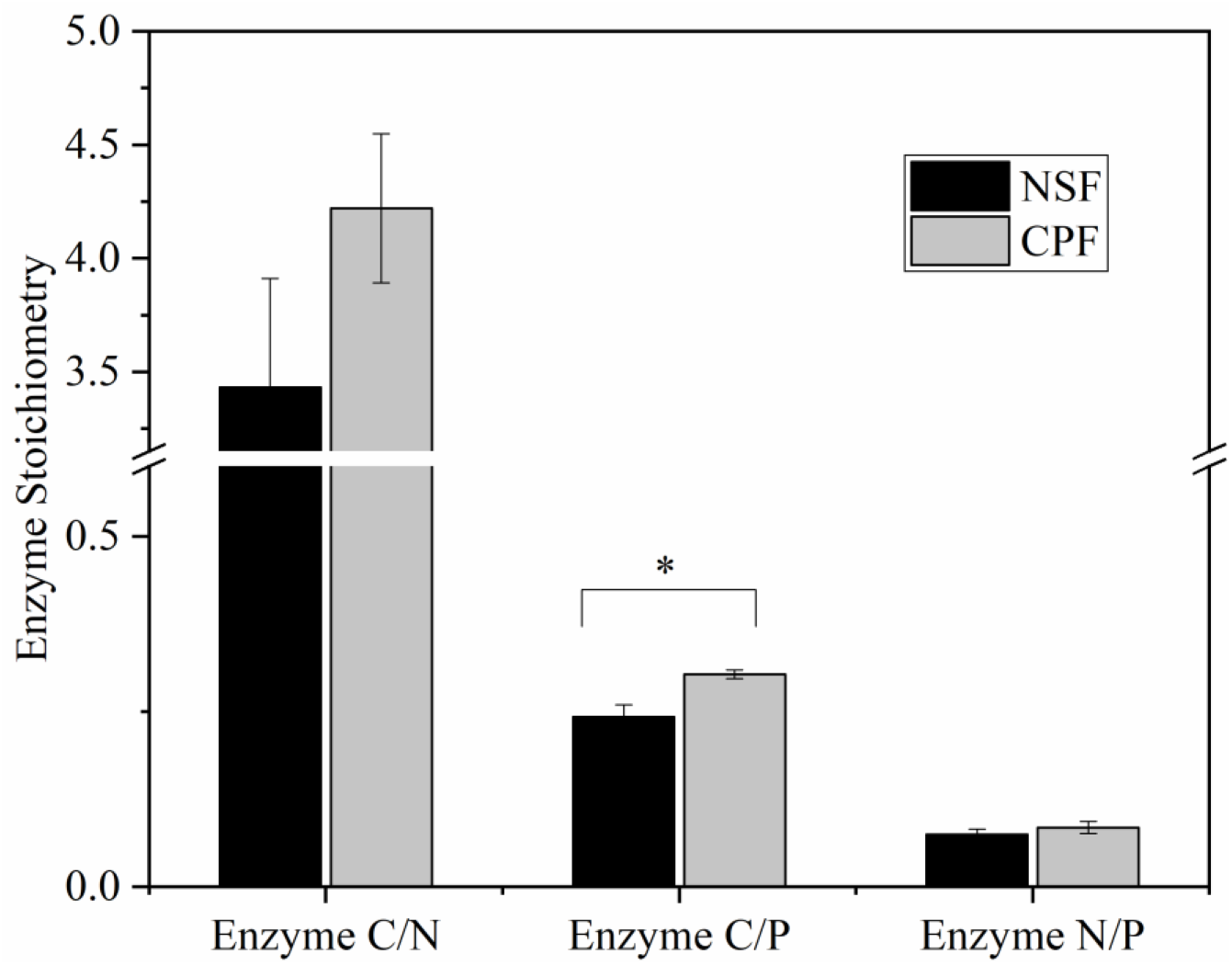
Comparisons of enzymatic stoichiometry at the 0-60 cm soil depth of the natural secondary forest (NSF) and Chinese fir plantation forest (CPF). All values are presented as mean ± standard error. “*” indicates a significance at a *p* <0.05 level.

The activity ratio of C/P in the CPF was significantly higher than that in the NSF (Fig. 3). The enzymatic C/P ratios at 10-20 and 40-60 cm in the CPF were 0.30 and 0.40 respectively, significantly greater than for the NSF. The highest C/P ratios were observed at 20-40 cm in the NSF (0.34) and 40-60 cm in the CPF (0.40) (Fig. 4). Enzymatic ratios of N/P at 0-10 and 10-20 cm in the NSF and the CPF were much lower than at 20-40 and 40-60 cm, while C/N in the CPF at 0-10 and 10-20 cm depths were significantly greater than the other two soil layers (Fig. 4).

**Fig. 4.**
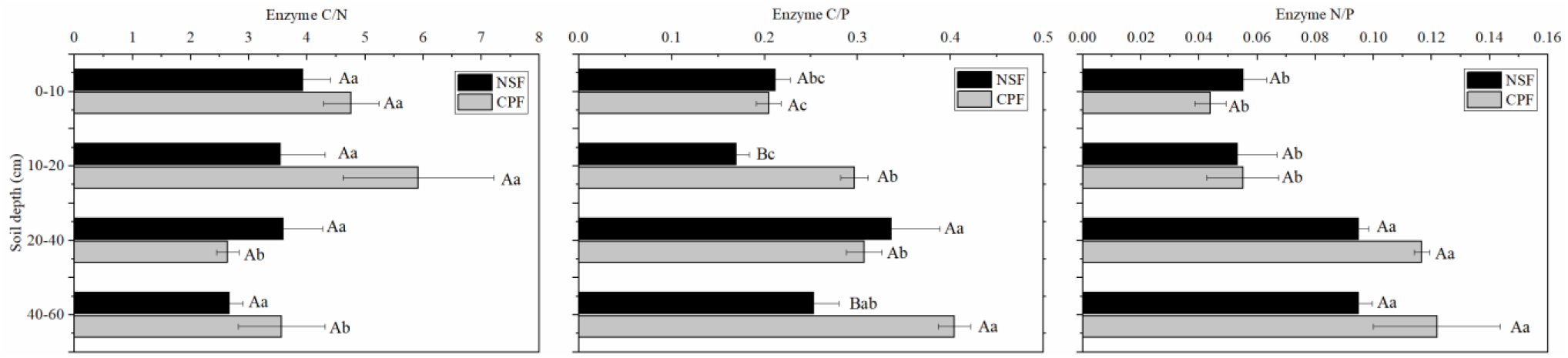
Variations of enzymatic stoichiometry at different soil depths of the natural secondary forest (NSF) and Chinese fir plantation forest (CPF). Capital letters mean the significant difference between the two forests at the same soil depth, and the different lower cases reflect the significant difference between soil depths within one forest. All values are presented as mean ± standard error with 3 replicates.

### 3.2 Vector analysis

Vector A (angle) in the CPF and NSF were greater than 45°in each layer that indicated microbial nutrient limitation in the two forest types was P limited at all soil depths and not N limited (Table 3). When combined with enzymatic stoichiometry scatter plots, our study indicated that all 4 soil layers were both C and P limited but not N limited (Table 3; S2 Fig.). The vector L (length) and A (angle) were not significantly different between NSF and CPF at all 4 sampling depths indicating the lack of differences in C- and P-limitation between NSF and CPF (Table 3).

**Table 3.** Vector analysis between the natural secondary forest (NSF) and Chinese fir plantation forest (CPF) at different soil depth.

| Depth<br>(cm) | Vector Length |  | Vector Angle |  |
| --- | --- | --- | --- | --- |
|  | NSF | CPF | NSF | CPF |
| 0-10 | 0.81 ±0.02Aa | 0.84 ±0.02Aa | 77.61 ±0.82Aab | 78.37 ±0.62Aa |
| 10-20 | 0.78 ±0.04Aa | 0.88 ±0.03Aa | 79.25 ±0.87Aa | 74.83 ±0.78Ab |
| 20-40 | 0.81 ±0.04Aa | 0.76 ±0.02Ab | 72.23 ±1.24Ac | 72.05 ±0.50Ac |
| 40-60 | 0.75 ±0.02Aa | 0.82 ±0.03Aab | 74.54 ±0.95Abc | 69.48 ±0.79Ad |

Values represented mean ± standard error (n=3). Capital letters show the significant difference between the two forests at the same soil depth, and the different lower cases reflect the significant difference between soil depths within one forest type.

Results of vector L demonstrated that there were no significant differences among the four soil layers in the NSF, while Vector L at 20-40 and 40-60 cm soil layers was notably lower than at 0-10 and 10-20 cm in CPF (Table 3). Although vector A varied significantly with depth in CPF, there was no coincident tendency for vector A in the NSF. In the NSF, the angles at each soil layer were greater than 45° and the largest and smallest angles occurred in the 10-20 and 20-40 cm soil depths, respectively.

### 3.3 Effects of soil properties on enzymatic activities and stoichiometries

The CCA indicated that the first two axes contained 69.4% of the total variance in soil enzymatic activities and the first and second axes explained 59.70 and 9.70% of the variance in the data, respectively (Fig. 5A). BG and C-acquiring activities were strongly negatively correlated with TN and TN/TP while CBH, NAG and BX activities were strongly positively correlated with most soil nutrients and nutrient stoichiometries. LAP activity and total N-acquiring activity were strongly negatively correlated with most soil nutrients and nutrient stoichiometries. The variations in soil enzymatic stoichiometry were well accounted for (71.2%) by soil properties and nutrient ratios (Fig. 5B). The enzyme C/P ratio was strongly negatively correlated with TN content and enzyme C/N was highly positively correlated with most soil nutrients and nutrient stoichiometries.

**Fig. 5.**
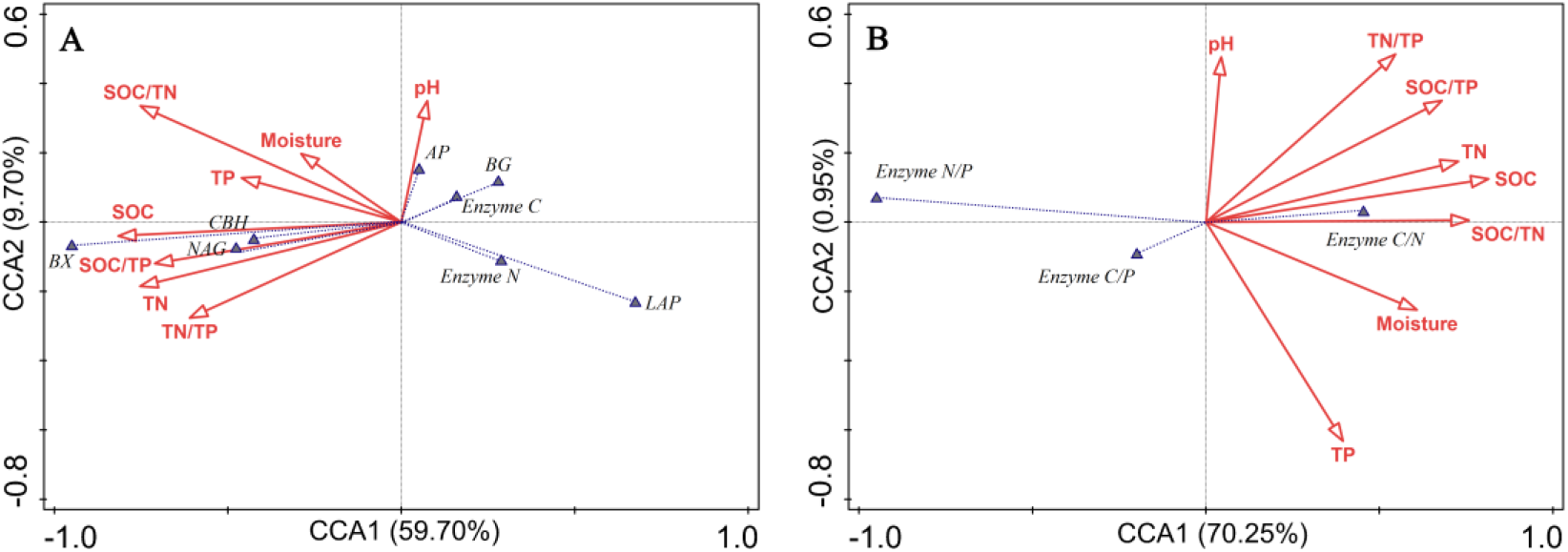
Canonical correspondence analysis (CCA) between environmental variables and (A) soil enzyme activities, and (B) soil enzyme stoichiometry. Moisture: soil moisture, SOC: soil organic carbon, TN: total nitrogen, TP: total phosphorus, BG: β-1,4-glucosidase, BX: β-xylosidase, CBH: β-D-cellobiosidase, NAG: β-1,4-Nacetylglucosaminidase, LAP: L-leucine aminopeptidase, AP: acid phosphatase. Enzyme C, Enzyme N and Enzyme P represent (BG + BX + CBH), (NAG + LAP) and AP, respectively. Enzyme C/N, C/P and N/P represent (BG + BX + CBH)/(NAG + LAP), (BG + CBH)/AP and (NAG + LAP)/AP, respectively.

## 4 Discussion

### 4.1 Spatial variations of C-, N- and P- acquiring enzyme activities

Generally, hydrolase activities decreased gradually with soil depth as has been observed previously [17, 35, 36]. We observed that C and P acquiring enzyme activities exhibited a decreasing trend with soil depth but not for all N- acquiring enzyme activities in the NSF. The largest fluctuations of enzyme activity occurred in 10-20 cm soil layer where C- acquiring enzyme activity in NSF and N- acquiring enzyme activity in both NSF and CPF decreased sharply. These results indicated that microbial metabolism was higher at 0-10 cm and can be explained by fine root biomass and microbes for these forests that was abundantly distributed in the top 0–10 cm soil layer [45]. Active microbial activities were present due to the rich and easily-degradable organic substances in the topsoil [46, 47]. No remarkable differences in C-acquiring enzyme activities such as BG were found between the NSF and CPF in the 0-10 cm soil layer although BX and CBH activities were significantly higher in the NSF at 0-10 cm (S1 Table). The CCA indicated that CBH and BX activities were strongly positively correlated with most soil nutrients (Fig. 5A) and this was consistent with SOC and TN levels that were higher in 0-10 cm in the NSF than in the CSF (Table 1). Although enzymes are substrate specific and individual enzyme activity may not reflect total soil nutritional status [48], many studies have utilized BG activity as the entire C acquiring enzyme because glucose released from cellulose is the most abundant monomer in soil [18]. Oligo- and monosaccharides that include glucose, xylose, arabinose, galactose, mannose and rhamnose are released from polysaccharide decomposition by microbial enzymes including cellulases, xylanases, glucosidases and chitinases. The released sugars are soluble and captured rapidly by microbes for metabolism and carbon storage [18]. Complex belowground biochemical processes indicate that using a single enzyme to describe the mechanism of microbial nutrient demands may be biased and using more enzymes in addition to BG should be studied to gain a true reflection of the sources and destinations of these sugars.

In the present study, significantly higher N-acquiring activity (NAG + LAP) was found at the surface soil in the NSF than in the CPF that was due to higher LAP activity (Fig. 2). LAP contributed almost 50% to the total N- acquiring enzyme activity (S1 Table). N-acquiring enzyme activity in the CPF did not decrease with soil depth and was associated with LAP activity (Fig. 2). The LAP distribution pattern was biphasic and increased after an initial decrease with increasing soil depth (S1 Table). This was consistent with previous studies indicating a major role for LAP in forest soil [6]. Additionally, NAG is often used as the indicator for N acquisition [35] and has been found to be positively correlated or no related to SOM [4, 17]. We found that NAG activity and the whole N-acquiring enzyme were strongly positively correlated with most soil nutrients and nutrient stoichiometries (Fig. 5A).

Previous studies have indicated that the Chinese fir mobilizes P for uptake through soil acidification and produce less acid phosphates [49, 50]. This results in high P demand for its rapid growth with lower AP activity, therefore, we found no differences in enzyme P activity at the whole soil profile in the CPF compared with the NSF (Fig. 2).

### 4.2 Enzymatic stoichiometry and microbial nutrient limitations

In the current study, both the scatter plot of enzymatic stoichiometry (S2 Fig.) and the vector analysis (Table 3) indicated that the two forests were commonly C and P limited at all soil depths and was consistent with karst ecosystems in southwest China [30]. Soil P is mostly unavailable to microbes such as bound to iron minerals or stabilized organic matter. The soil pH can alter complexed P and make P more available to microbes [51]. We indeed found the CPF had a higher topsoil pH than the NSF, however, vector A between NSF and CPF in the topsoil did not show a significant difference. Furthermore, the lower enzymatic N/P and C/P ratios were both <1 at all depths indicating P limitation in the the two forests [4, 52]. A gradual decrease of vector A in the CPF indicated that microbial P limitation was gradually reduced along soil depth. While vector A fluctuated according to soil depth in the NSF, the highest microbial P limitation was at 10-20 cm and lowest at 20-40 cm (Table 3). Results of mean vector A indicated that microbial P limitation were higher in the 0-20 cm soil layer than in the 20-60 cm in both forests, and microbial P limitation in the 20-60 cm soil layer in the NSF was significantly higher than that in the CPF which could be because Chinese fir is shallow rooted species (S4 Fig.). Previous studies have revealed that P restrictions are quite common and our study is in agreement with this in mid-subtropical forests [5, 53, 54].

Subtropical ecosystems are N-rich relative to other plant nutrients and N was not limiting with respect to microbial growth demand as evidenced from the enzymatic stoichiometry results [55–57]. That is, the C/N ratio may not reach the critical ratio of the two elements for microbial growth. According to the N-mining theory, some microbes can use labile C to decompose recalcitrant organic matter to obtain N, which implies that microbial N-acquisition can be alleviated by C-acquisition [58]. The sharp increases of the enzymatic N/P ratio and significantly lower of vector L below 20 cm indicated a greater N demand of microbes in deep soils for both forests and were consistent with results in a subtropical mixed forest soil (Fig. 4; S4 Fig.) [59].

Although the differences of enzyme C/N and C/P ratios between the NSF and CPF at all soil layers remained unchanged when using single BG as whole C acquisition enzyme, the enzyme C/P ratio at NSF displayed a different distribution trend along the soil depths. The enzyme C/P ratio between 10-20 and 40-60 cm in the NSF changed from significant to insignificant after using single BG as the sole indicator of C acquisition (Fig. 4; S3 Fig.). This underestimated the microbial C demand of NSF at the 40-60 cm soil depth. On the other hand, vector A in NSF at 40-60 cm depth became significantly higher than that in CPF when using only BG (S5 Table). Furthermore, vector A between 10-20 and 40-60 cm in the NSF changed from significant to insignificant. These two alterations demonstrated that microbial P limitation in the NSF at 40-60 cm would be overestimated when using only BG as the sole C acquisition enzyme. In conclusion, utilizing single BG as whole C acquisition enzyme may miscalculate microbial C and P demand in deep soils.

Although previous studies and our study indicated that enzymatic stoichiometry can be used to identify microbial nutrient limitation, enzymes are snapshot proxies for a complex plastic expression of microbial cellular metabolism, and the assayed enzyme activity might not fully reflect actual microbial metabolism, because the enzyme activities might also come from enzymes stabilized in the soil matrix [48]. Moreover, the stoichiometry could well indicate the quantitative relationships with elements in soil and enzyme. We observed that the strong relationships between soil nutrient stoichiometry and enzyme activities together with enzymatic stoichiometry, which might imply the complicated responses of microbial communities to altered resources in the two forests, and the fluctuation of microbial nutrient limitation with time (Fig. 5). To more accurately determine the microbial nutrient limitation in mid-subtropics, long-term analysis of enzymatic stoichiometry or joint observation with other methods is necessary.

East Asian monsoon subtropical forests have one of the highest carbon uptakes of forests worldwide and are the result of young stand ages and high N deposition [60]. Nevertheless, soil total C levels have increased significantly and soil total P concentration decreased significantly across all soil depths in subtropical China from 1955 to 2016 [56]. To the best of our knowledge, this study represents novel findings in examining microbial nutrient demands along a whole soil profile in mid-subtropical forests and can contribute to informational references for forest management in the subtropics.

### 4.3 Soil stoichiometry and microbial C and P limitations

Ecological stoichiometry refers to the balance between energy and various chemical elements by element ratios [11, 61], and soil stoichiometry (soil C:N:P ratio) have long been considered as useful indicators of dynamics of soil fertility [62, 63]. It is reported that ecological stoichiometric theory and the metabolic theory of ecology can be related to enzymatic C:N:P stoichiometry via the threshold elemental ratio concept, and enzymatic stoichiometry can be used to determine energetic and nutrient constraints on microbial community metabolism [11]. Soil stoichiometry may cause changes in microbial interactions and community dynamics that can lead to feedbacks in nutrient availability [53]. However, the roles of soil stoichiometries in microbial nutrient limitation have not been active areas of research especially for forest ecosystems [64]. Our study indicated that microbial C and P limitations were significantly affected by soil stoichiometry across the whole soil profile (S4 Table). The calculations suggested that soil C:P ratios > 10 and C:N ratios > 8 can also be indicators of microbial C and P limitations in the study (Table 3, S5 Table). The reasons could attribute to microorganisms adapting to the altered resources in the NSF and CPF, and holding elemental stoichiometric balance and homeostasis [53]. Therefore, our findings indicated that soil C, N and P stoichiometry can significantly regulate microbial community metabolism and that the stoichiometry can provide references for understanding the mechanisms of nutrient cycling through determination of microbial nutrient metabolic limitations.

## 5 Conclusion

C-, N- and P-acquiring enzyme activities decreased with increasing soil depth but there were few differences in enzyme activities along the whole soil profile between the subtropical secondary forests and Chinese fir plantations. Moreover, the results of enzyme stoichiometry indicated higher N demand by microbes in soils below 20 cm in both forests. Microbial C and P but not N limitations were observed at all soil depths in both forests. The insignificant differences for microbial C, N and P limitation between the two forests at most soil depths implied complex belowground plant-microbe interactions. The microbial C and P limitations were remarkably correlated with soil C N and P stoichiometry in the whole soil profile, indicating that the imbalance of stoichiometric ratio might be important factor affecting microbial metabolism in terrestrial ecosystems. Utilizing single BG as the sole C acquisition enzyme could result in miscalculations of microbial C and P demands at deeper soil layers in the study. The present study gives a new insight into subtropical forests in examining microbial nutrient demands in both top and deep soils, and provides essential information for subtropical tree plantation and natural forest management.

## 6 Acknowledgments

We thank Daryl Moorhead for his valuable suggestions in preparing the manuscript, and we also thank Teng-Chiu Lin and the anonymous reviewers together with the editors of the journal who provided comments and suggestions on the manuscript.

## Appendix A supplementary

