## Supplemental figs and tables for "Enzyme stoichiometry indicates the variation of microbial nutrient requirements at different soil depths in subtropical forests"

### 1 Supporting Information

**S1 Table.** **Individual enzyme activity in the natural secondary forest (NSF) and Chinese fir plantation forest (CPF) at different soil depths**

| Depth | BX  (nmol g^-1^ dry soil h^-1^) | |  | CBH  (nmol g^-1^ dry soil h^-1^) | |  | BG  (nmol g^-1^ dry soil h^-1^) | |  | NAG  (nmol g^-1^ dry soil h^-1^) | |  | LAP  (nmol g^-1^ dry soil h^-1^) | |
| --- | --- | --- | --- | --- | --- | --- | --- | --- | --- | --- | --- | --- | --- | --- |
| (cm) | NSF | CPF |  | NSF | CPF |  | NSF | CPF |  | NSF | CPF |  | NSF | CPF |
| 0-10 | **149.9±6.1Aa** | **92.4±8.4Ba** |  | **43.9±2.9Aa** | **31.8±2.2Ba** |  | 360.0±53.9Aa | 404.8±61.5Aa |  | 75.6±4.2Aa | 55.1±4.5Aa |  | **65.5±2.6Aa** | **57.0± 3.0Bab** |
| 10-20 | 74.9± 13.5Ab | 78.1± 19.9Aa |  | 24.3±0.9Ab | 19.9± 2.0Ab |  | 227.7±60.0Aab | 302.5±6.7Aa |  | 59.3± 7.3Ab | 39.0± 4.8Aab |  | 34.0±6.6Ab | 33.5±8.4Ab |
| 20-40 | 46.1 ±11.3Abc | 34.5±13.6Ab |  | 15.6±2.7Ac | 18.7± 2.5Ab |  | 277.1±60.9Aab | 191.5± 5.6Ab |  | 36.4±2.5Ac | 31.9 ±3.8Acdc |  | 56.8±3.4Aa | 60.6± 3.0Aa |
| 40-60 | 21.9± 3.3Ac | 14.5±4.3Ab |  | 10.8±2.0Ac | 13.7± 1.6Ab |  | 153.0±15.0Ab | 175.6±13.6Ab |  | 18.3±1.7Ad | 17.7± 4.6Ad |  | 52.1±8.1Aab | 42.8±11.7Aab |

Capital letters mean the significant difference between the two forests at the same soil depth, and the different lower cases reflect the significant difference between soil depths within one forest. Values were mean ± standard error (n=3). BX:β-xylosidase, CBH: β-D-cellobiosidase, BG: β-1,4-glucosidase, NAG: β-1,4-N-acetylglucosaminidase, LAP: L-leucine aminopeptidase.

**S2 Table**. **Spearman correlation coefficients (ρ) relating extracellular enzyme activities with soil chemical properties and nutrient stoichiometry**.

| Enzyme activity  (nmol g soil^-1^ h^-1^) | pH | Moisture  (%) | SOC  (g kg^-1^) | TN  (g kg^-1^) | TP  (g kg^-1^) | SOC/TN | SOC/TP | TN/TP |
| --- | --- | --- | --- | --- | --- | --- | --- | --- |
| BG+BX+CBH | -0.364 | 0.682** | 0.825** | 0.827** | 0.482* | 0.625** | 0.689** | 0.580** |
| LAP+NAG | -0.352 | 0.559** | 0.788** | 0.808** | 0.267 | 0.423* | 0.694** | 0.688** |
| AP | -0.201 | 0.695** | 0.908** | 0.864** | 0.417* | 0.671** | 0.781** | 0.656** |
| BX | -0.241 | 0.517** | 0.917** | 0.892** | 0.435* | 0.667** | 0.790** | 0.699** |
| CBH | -0.094 | 0.543** | 0.864** | 0.832** | 0.470* | 0.671** | 0.755** | 0.646** |
| BG | -0.380 | 0.688** | 0.712** | 0.727** | 0.474* | 0.556** | 0.585** | 0.474* |
| NAG | -0.188 | 0.541** | 0.879** | 0.865** | 0.336 | 0.621** | 0.781** | 0.708** |
| LAP | -0.411* | 0.249 | 0.135 | 0.193 | -0.004 | -0.153 | 0.114 | 0.217 |

**Note:** Moisture: soil moisture, SOC: soil organic carbon, TN: total nitrogen, TP: total phosphorus, BG: β-1,4-glucosidase, CBH: β-D-cellobiosidase, BX: β-xylosidase, NAG: β-1,4-N-acetylglucosaminidase, LAP: L-leucine aminopeptidase, AP: acid phosphatase. Correlations were considered significant (*) at *p* < 0.05 (two-tailed) and highly significant (**) at *p* < 0.01 (two-tailed).

**S3 Table**. **Spearman correlation coefficients (ρ) relating enzyme stoichiometry with soil chemical properties and nutrient stoichiometry**.

| Enzyme stoichiometry | pH | Moisture  (%) | SOC  (g kg^-1^) | TN  (g kg^-1^) | TP  (g kg^-1^) | SOC/TN | SOC/TP | TN/TP |
| --- | --- | --- | --- | --- | --- | --- | --- | --- |
| (BG+BX+CBH)/(NAG+LAP) | -0.210 | 0.505* | 0.543** | 0.530** | 0.512* | 0.531** | 0.415* | 0.281 |
| (BG+BX+CBH)/ AP | -0.172 | -0.465* | -0.636** | -0.563** | -0.039 | -0.456* | -0.610** | -0.511* |
| (NAG+LAP)/ AP | 0.008 | -0.626** | -0.777** | -0.728** | -0.380 | -0.641** | -0.668** | -0.515* |

Moisture: soil moisture, SOC: soil organic carbon, TN: soil total nitrogen, TP: soil total phosphorus, BG: β-1,4-glucosidase, CBH: β-D-cellobiosidase, BX: β-xylosidase, NAG: β-1,4-N-acetylglucosaminidase, LAP: L-leucine aminopeptidase, AP: alkaline phosphatase. * Correlation is signiﬁcant at *p* < 0.05 (two-tailed); ** Correlation is highly signiﬁcant at *p* < 0.01 (two-tailed).

**S4 Table**. **Spearman correlation coefficients (ρ) relating vector length and angle with soil chemical properties and nutrient stoichiometry**.

| Vector characteristics | pH | Moisture  (%) | SOC  (g kg^-1^) | TN  (g kg^-1^) | TP  (g kg^-1^) | SOC/TN | SOC/TP | TN/TP |
| --- | --- | --- | --- | --- | --- | --- | --- | --- |
| Length | -0.228 | 0.414* | 0.419* | 0.425* | 0.488* | 0.420* | 0.301 | 0.195 |
| Angle | 0.131 | 0.564** | 0.754** | 0.685** | 0.150 | 0.580** | 0.711** | 0.583** |

Moisture: soil moisture, SOC: soil organic carbon, TN: soil total nitrogen, TP: soil total phosphorus. * Correlation is signiﬁcant at *p* < 0.05 (two-tailed); ** Correlation is highly signiﬁcant at *p*< 0.01 (two-tailed).

**S5 Table.** **Vector analysis, usingβ-1,4-glucosidase (BG) as single C acquiring enzyme, between the natural secondary forest (NSF) and the Chinese fir plantation forest (CPF) at different soil depths**

| Depth | Vector L | |  | Vector A | |
| --- | --- | --- | --- | --- | --- |
| (cm) | NSF | CPF |  | NSF | CPF |
| 0-10 | 0.72±0.03Aa | 0.79±0.03Aa |  | 80.46±0.57Aab | 80.22±0.65Aa |
| 10-20 | 0.69±0.07Aa | 0.83±0.02Aa |  | 81.45±0.50Aa | 77.14±1.04Ab |
| 20-40 | 0.77±0.04Aa | 0.70±0.003Ab |  | 73.90±1.11Ab | 73.92±0.19Ac |
| 40-60 | 0.71 ±0.02Aa | 0.79±0.04Aab |  | 75.96±0.87Aab | 70.79±0.95Bd |

Values represented mean ± standard error (n=3). Capital letters show the significant difference between the two forests stand at the same soil depth, and the different lower cases reflect the significant difference within one forest stand at four soil depths.

**S1 Fig (A) the natural secondary forest (NSF) and (B) the Chinese fir plantation forest (CPF)**

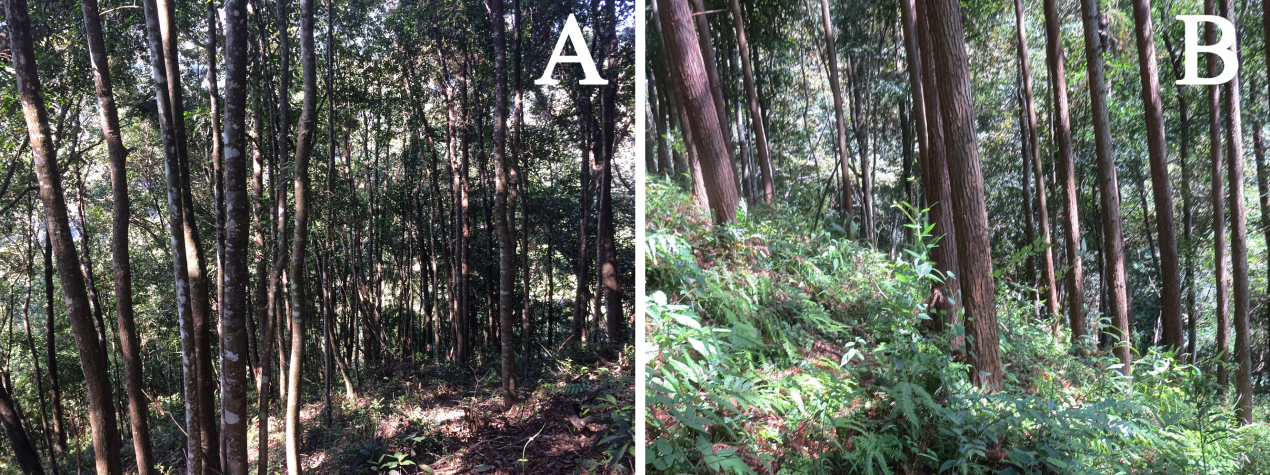

**S2 Fig. Scatter plots of soil enzymatic stoichiometry. Red and blue colors represent the natural secondary forest (NSF) and Chinese fir plantation forest (CPF) respectively**. Different symbols represent different soil depth.

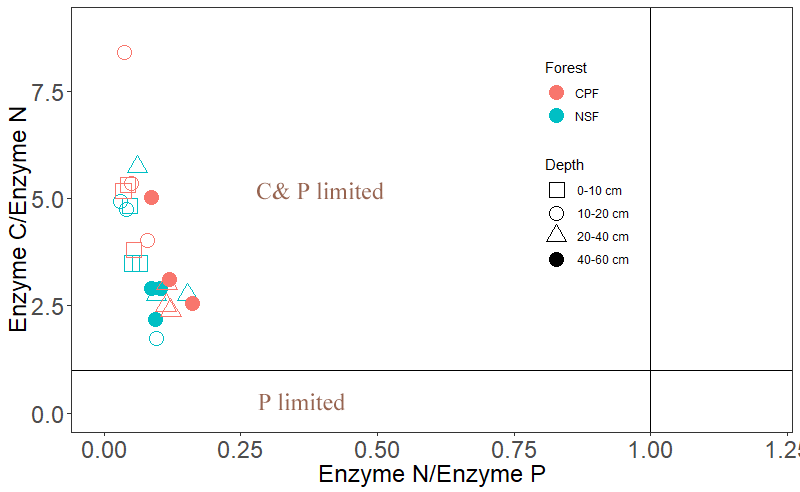

**S3 Fig. Enzyme stoichiometry of BG/AP and BG/(NAG+LAP) at different soil depths**. Capital letters mean the significant difference between the natural secondary forest (NSF) and Chinese fir plantation forest (CPF) at the same soil depth, and the different lower cases reflect the significant difference between soil depths within one forest. Comparison between the two forests at the same soil depth was performed by paired-sample t-test. One-way analysis of variance (ANOVA) with LSD multiple comparisons among four soil depths was conducted at the same forest type. All values are presented as mean ± standard error (n=3). BG, β-1,4-glucosidase; AP, acid phosphatase; NAG, β-1,4-N-acetylglucosaminidase and LAP, leucine aminopeptidase.

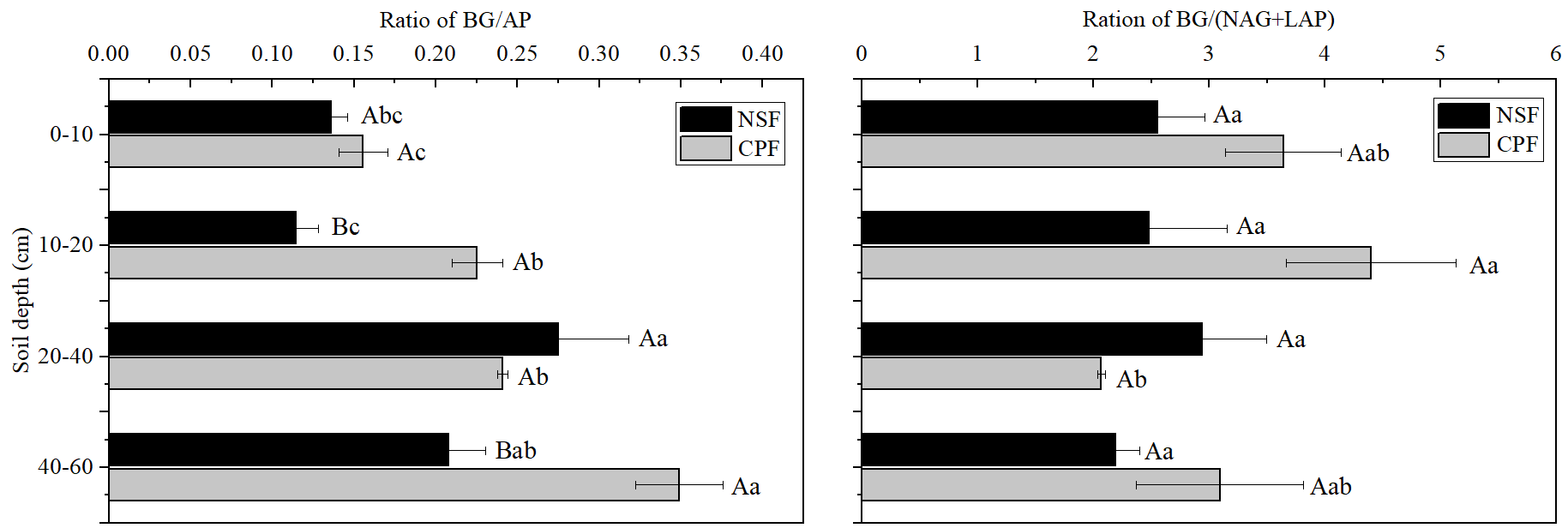

**S4 Fig.** **Vector analysis between the natural secondary forest (NSF) and Chinese fir plantation forest (CPF) at different soil depths**. Values represented mean ± standard error (n=3). Different capital letters show the significant difference between the two forests stand at the same soil depth, and the different lower cases reflect the significant difference within the same forest type at two soil depths.

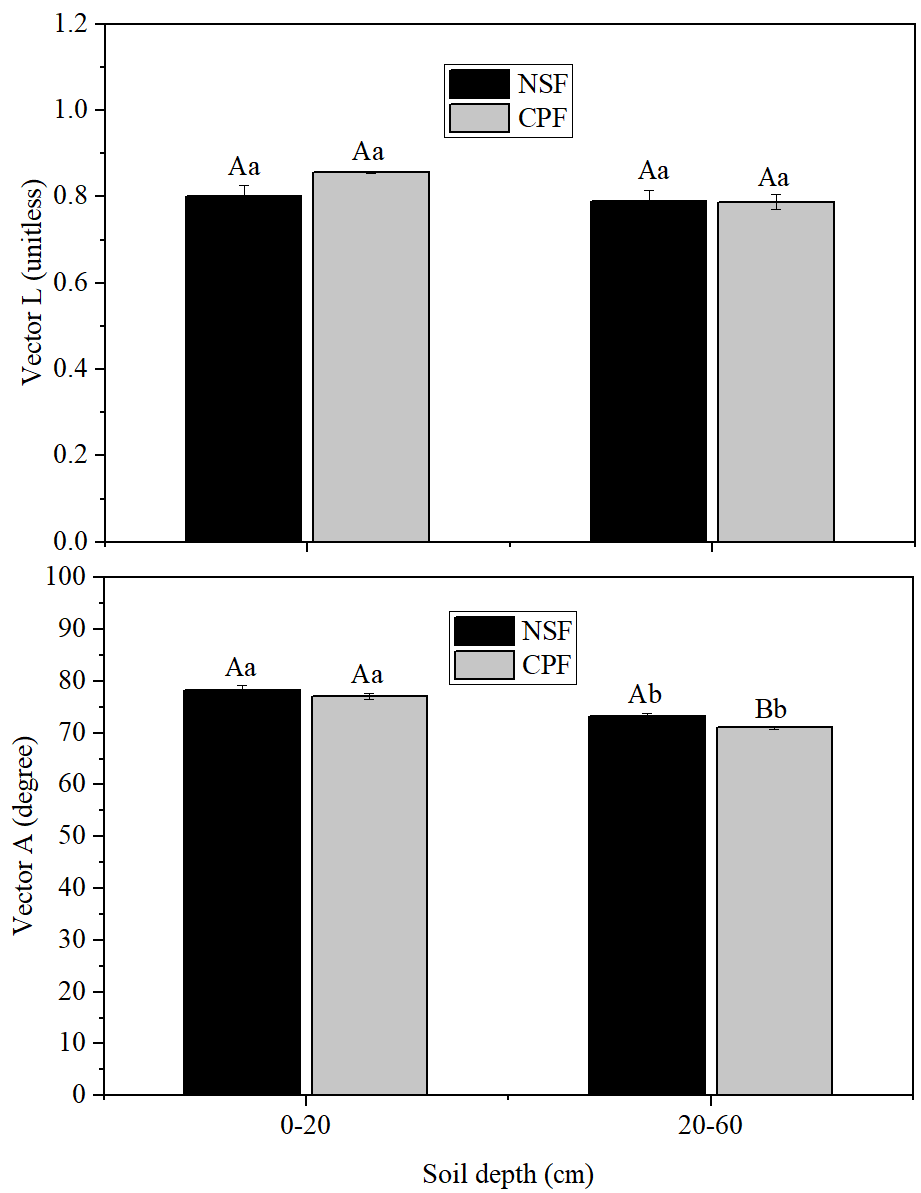
